# Mitochondrial-nuclear heme trafficking is regulated by GTPases that control mitochondrial dynamics

**DOI:** 10.1101/539254

**Authors:** Osiris Martinez-Guzman, Jonathan V. Dietz, Iryna Bohovych, Amy E. Medlock, Oleh Khalimonchuk, Amit R. Reddi

## Abstract

Heme is an essential cofactor and signaling molecule. All heme-dependent processes require that heme is trafficked from its site of synthesis in the mitochondria to hemoproteins in virtually every subcellular compartment. However, the mechanisms governing the mobilization of heme out of the mitochondria, and the spatio-temporal dynamics of these processes, are poorly understood. To address this, we developed a pulse-chase assay in which, upon the initiation of heme synthesis, heme mobilization into the mitochondrial matrix, cytosol and nucleus is monitored using fluorescent heme sensors. Surprisingly, we found that heme trafficking to the nucleus occurs at a faster rate than to the matrix or cytosol. Further, we demonstrate that GTPases in control of mitochondrial fusion, Mgm1, and fission, Dnm1, are positive and negative regulators of mitochondrial-nuclear heme trafficking, respectively. We also find that heme controls mitochondrial network morphology. Altogether, our results indicate that mitochondrial dynamics and heme trafficking are integrally coupled.

## Introduction

Heme (iron protoporphyrin IX) is an essential but inherently cytotoxic metallocofactor and signaling molecule^1,2^. As a cofactor, heme facilitates diverse processes that span electron transfer, chemical catalysis, and gas synthesis, storage and transport^1,2^. As a signaling molecule, heme regulation of an array of proteins, including transcription factors^3–6^, kinases^7,8^, ion channels^9^, micro RNA processing proteins^10^, collectively control pathways spanning iron homeostasis, oxygen sensing, the oxidative stress response, mitochondrial respiration and biogenesis, mitophagy, apoptosis, circadian rhythms, cell cycle progression, proliferation, and protein translation and degradation^1,2^. Although essential for life, heme may also act as a toxin, necessitating that cells carefully handle this compound. The hydrophobicity and redox activity of heme causes it to disrupt membrane structure, become mis-associated with certain proteins, and deleteriously oxidize various biomolecules^11,12^. Indeed, a number of diseases are associated with defects in heme management, including certain cancers^4^, cardiovascular disease^13^, aging and age-related neurodegenerative diseases^14–16^, porphyrias^17^, and anemias^18^. Despite the tremendous importance of heme in physiology, the cellular and molecular mechanisms that govern the *safe* assimilation of heme into metabolism remain poorly understood.

All heme dependent processes require that heme is mobilized from its site of synthesis on the matrix side of the mitochondrial inner membrane (IM) to heme proteins present in virtually every subcellular compartment^1,2^. However, the specific factors that govern the mobilization of heme out of the mitochondria, heme distribution to a multitude of locales, and heme insertion into target hemoproteins are not well-understood^1,2^. Our approach to better understand heme mobilization and utilization is to directly probe bioavailable or kinetically labile heme (LH) relevant for heme trafficking and signaling with fluorescent heme imaging agents^19^. Towards this end, we recently deployed a class of genetically encoded ratiometric fluorescent <u>h</u>eme <u>s</u>ensors (HS1) to the model unicellular eukaryote, *Saccharomyces cerevisiae*, and determined the subcellular distribution of LH, discovered new heme homeostatic factors, *e.g.* glyceraldehyde phosphate dehydrogenase (GAPDH), and signaling molecules that can mobilize LH, *e.g.* nitric oxide (NO)^19^. Altogether, the integration of heme imaging technologies with molecular genetic, cell biological, and biochemical approaches has revealed fundamental aspects of heme trafficking and dynamics, thereby providing fresh insight into the cellular management of this essential nutrient^1^.

In this contribution, we describe a new pulse-chase assay in which the spatio-temporal dynamics of heme mobilization from the mitochondrial IM to the matrix, cytosol, and nucleus is monitored with fluorescent heme sensors. Surprisingly, we find that heme transport rates from the matrix side of the IM to different cellular locales are similar, with heme transport rates to the nucleus being faster than to the mitochondrial matrix or cytosol. These data indicate that heme is distributed from the mitochondrial IM to other locales via multiple parallel pathways rather than sequentially. Further, we find that GTPases that control mitochondrial fusion and fission, Mgm1 and Dnm1, respectively, but not mitochondrial dynamics *per se*, are key regulators of heme trafficking from the mitochondrial IM. In particular, we find that Mgm1 is a positive regulator of mitochondrial-nuclear heme transport whereas Dnm1 is a negative regulator of heme trafficking to the matrix, cytosol, and nucleus. Further, we find that heme itself regulates mitochondrial network morphology. In total, the development of a new heme trafficking dynamics assay coupled with molecular genetics approaches have revealed heretofore unknown mechanisms of inter-compartmental heme trafficking.

## Results

### Inter-compartmental heme transport kinetics

In order to probe inter-compartmental heme trafficking in yeast, we developed a pulse-chase assay in which, upon the initiation of heme synthesis, heme mobilization into the mitochondrial matrix, cytosol and nucleus is monitored using genetically encoded ratiometric fluorescent <u>h</u>eme <u>s</u>ensors (HS1)^19^. HS1 is a tri-domain fusion protein consisting of a heme-binding moiety, the His/Met coordinating 4-alpha-helical bundle hemoprotein cytochrome *b*_562_ (Cyt *b*_562_), fused to a pair of fluorescent proteins, eGFP and mKATE2, that exhibit heme-sensitive and –insensitive fluorescence, respectively (**Fig. 1a**). Heme binding to the Cyt *b*_562_ domain results in the quenching of eGFP fluorescence via resonance energy transfer but has little effect on mKATE2 fluorescence. Thus, the ratio of eGFP fluorescence (ex: 488 nm, em: 510 nm) to mKATE2 fluorescence (ex: 588 nm, em: 620 nm) reports cellular heme independently of sensor concentration, with the eGFP/mKATE2 ratio inversely correlating with heme binding to the sensor. Throughout this study, unless otherwise noted, the sensor is expressed on a centromeric plasmid and the *GPD* promoter (p*GPD*) is used to drive HS1 expression. HS1 is targeted to the mitochondria and nucleus by appending N-terminal COX4 mitochondrial matrix or C-terminal SV40 nuclear localization sequences, respectively^19^ (**Fig. 1b and Fig. S1**).

**Figure 1.**
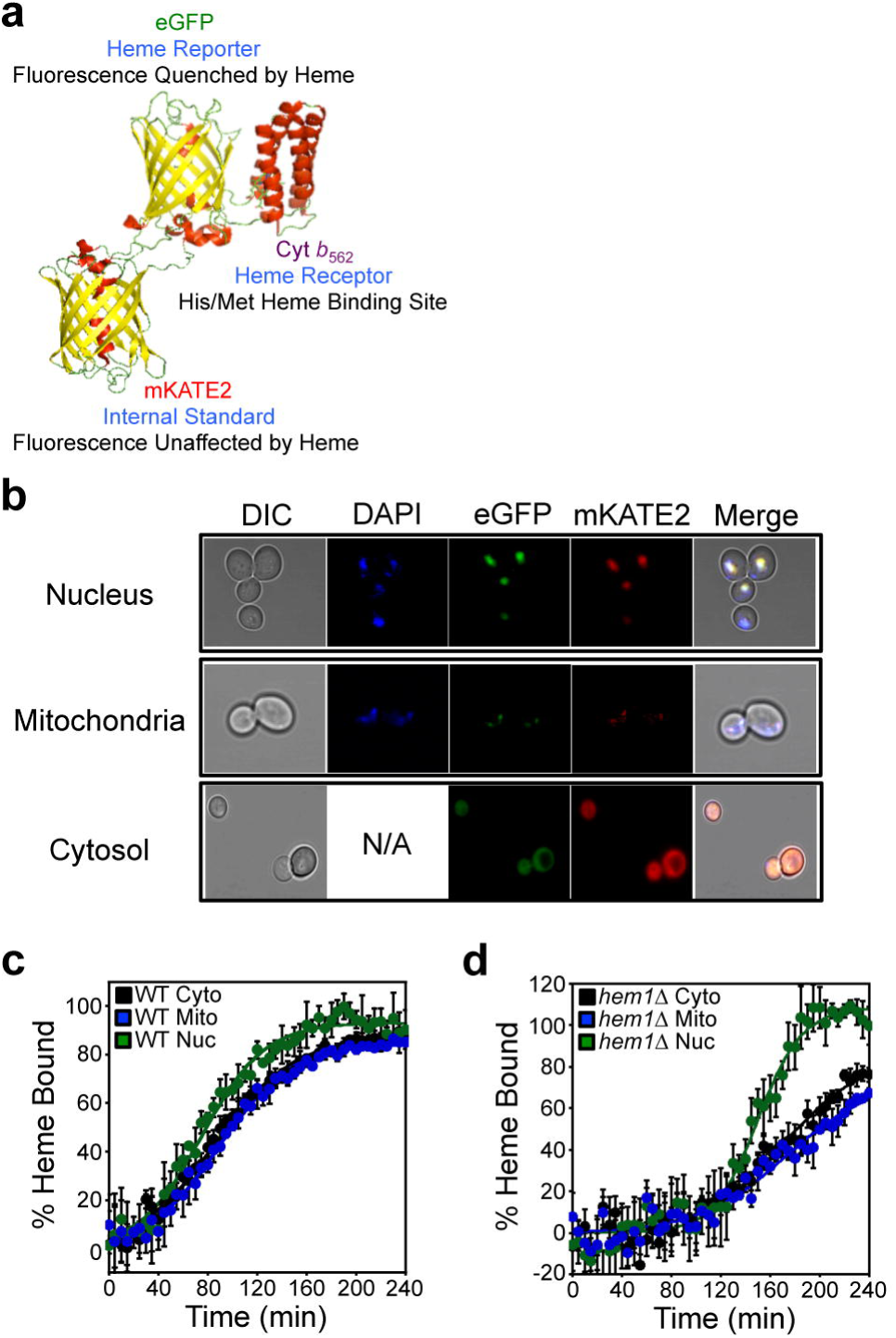
Inter-compartmental heme trafficking dynamics as measured by heme sensor HS1. (**a**) Molecular model and design principles of the heme sensor, HS1. Model derived from the X-ray structures of mKATE (PDB: 3BXB) and CG6 (PDB: 3U8P). (**b**) Validation of the sub-cellular localization of cytosolic, nuclear, and mitochondrial HS1 by laser scanning confocal microscopy. (**c**) WT cells expressing HS1 in the cytosol (black), nucleus (green), or mitochondria (blue) were depleted of heme using 500 μM succinylacetone (SA), the heme biosynthetic inhibitor and, upon the re-initiation of heme synthesis, the rates of heme trafficking to the indicated subcellular locations were monitored by measuring the fractional saturation of HS1 over time. The data are fit to equation 1 and the raw sensor fluorescence ratios giving rise to this data are provided in Fig. S2. (**d**) *hem1*∆ cells expressing HS1 in the cytosol (black), nucleus (green), or mitochondria (blue) were pulsed with a bolus of ALA to initiate heme synthesis and the rates of heme trafficking to the indicated subcellular locations were monitored by measuring the fractional saturation of HS1 over time. The heme trafficking kinetics data represent the mean ± SD of independent triplicate cultures. The kinetic parameters derived from the fits to the data are indicated in **Table 1**.

The fractional heme saturation of the sensor can be determined using previously established sensor calibration protocols^19^. The % of sensor bound to heme, % Bound, involves determining the sensor eGFP/mKATE2 fluorescence ratio (*R*) under a given test condition relative to the eGFP/mKATE2 fluorescence ratio when the sensor is 100% (*R*_max_) or 0% (*R*_min_) bound to heme as described in equation 1 and discussed in greater detail in the **Materials and Methods** section^19^.

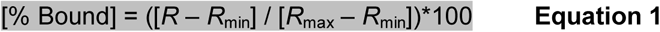

**Table 1.**
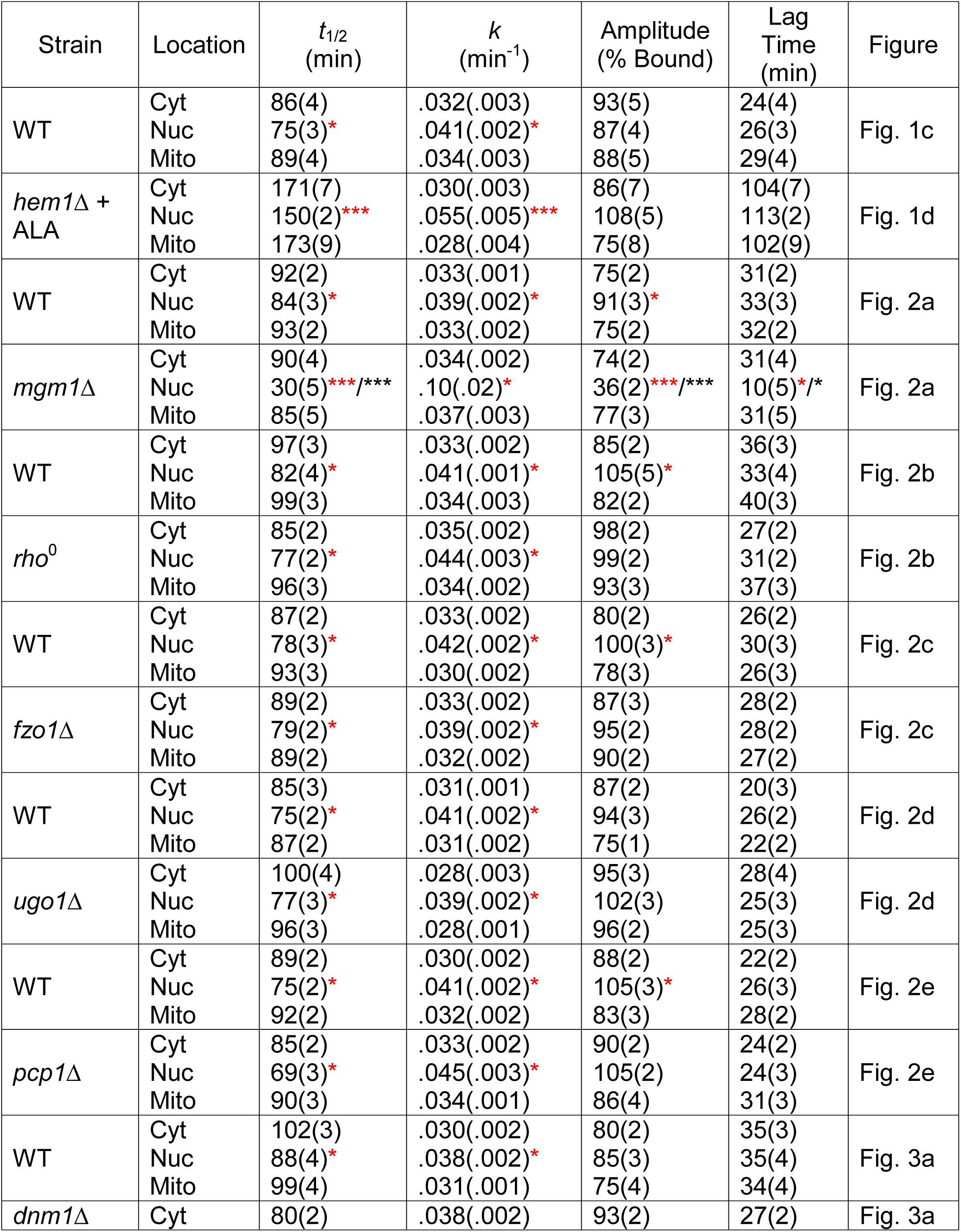
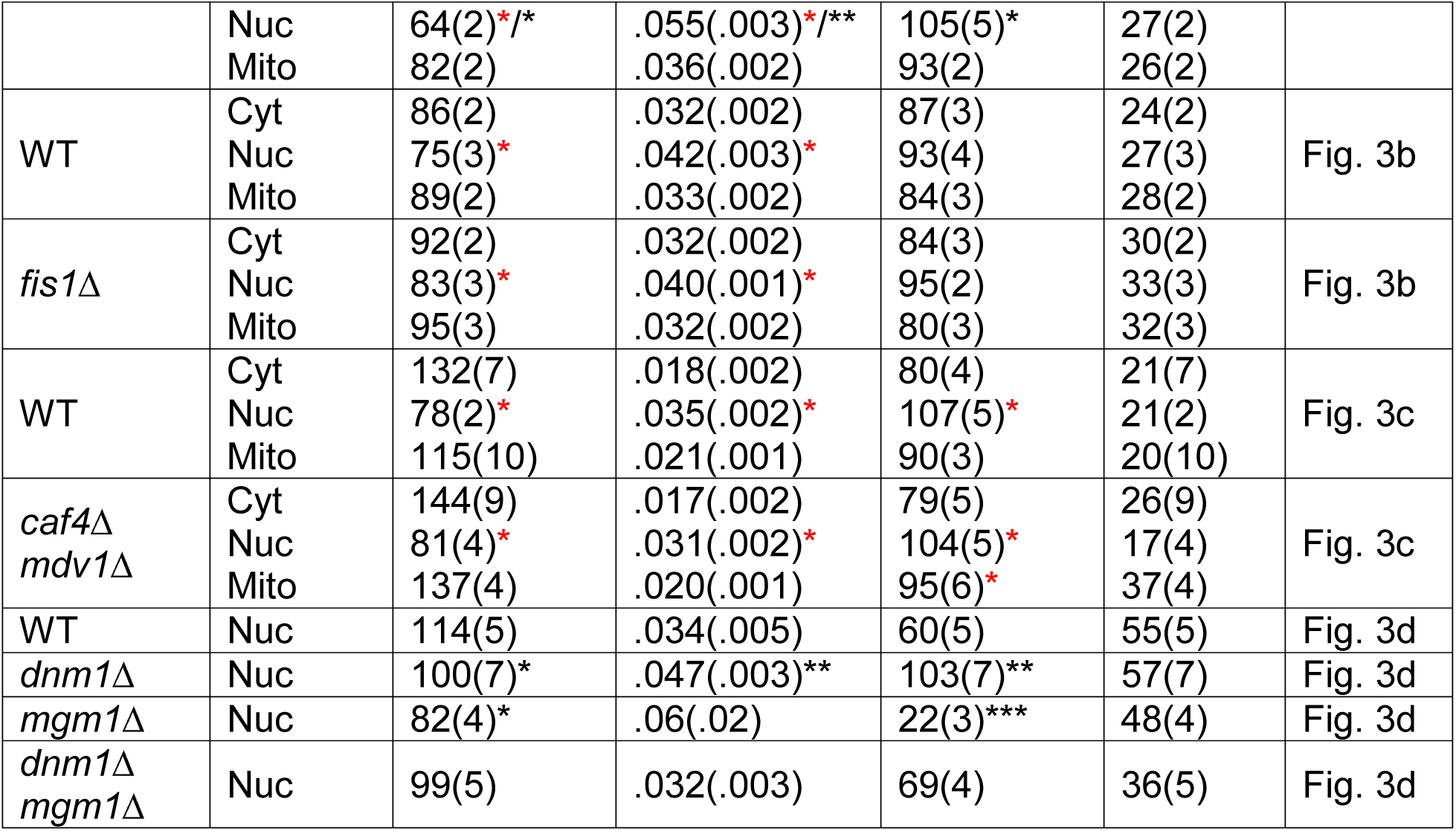
Kinetic parameters derived from fits to the heme trafficking data using Equation 2. The values indicated represent the mean ± SD of independent triplicate samples. *p<0.05, **p<0.01, ***p<0.001 by Student’s *t*-test. Red asterisks indicate statistical significance relative to the cytosol within a given strain. Black asterisks indicate statistical significance relative to the WT strain of a given compartment.

*R*_min_ is determined by measuring the HS1 eGFP/mKATE2 ratio in parallel cultures that are conditioned with succinylacetone (SA), which inhibits the second enzyme in the heme biosynthetic pathway, 5-aminolevulinic acid (ALA) dehydratase (ALAD)^20^, and *R*_max_ can be determined by permeabilizing cells and adding an excess of heme to saturate the sensor^19^. Given HS1 is quantitatively saturated with heme in the cytosol, nucleus, and mitochondria of WT yeast, we typically determine *R*_max_ by measuring the HS1 eGFP/mKATE2 ratio in parallel WT cultures grown without SA^19^.

Inter-compartmental heme trafficking rates are monitored by: **a.** inhibiting heme synthesis with 500 μM SA for ~16 hours in sensor expressing cells; **b.** removing the block in heme synthesis by re-suspending cells into media lacking SA; and **c.** monitoring the time-dependent change in the eGFP/mKATE2 ratio of HS1 localized to different locations upon the re-initiation of heme synthesis (**Fig. 1c and S2**). Parallel cultures of cells continually maintained in media with and without SA give HS1 *R*_min_ and *R*_max_ values, respectively (**Fig. S2**), which are used to calculate HS1 heme loading (% Bound) (**Fig. 1c**).

In order to confirm the SA-mediated block in heme synthesis, we determined that SA-conditioned WT cells exhibit intracellular heme concentrations^21^ (**Fig. S3a**) and HS1 eGFP/mKATE2 ratios^19^ (**Fig. S3b**) similar to *hem1*∆ cells, which cannot make heme due to the deletion of the first enzyme in the heme biosynthetic pathway, 5-aminolevulinic acid (5-ALA) synthase^19^. SA pre-conditioned cells shifted to media lacking SA exhibit a time-dependent decrease in HS1 eGFP/mKATE2 ratio (**Fig. S2**) in the cytosol, nucleus, and mitochondria, which is characteristic of increased heme loading of the sensor due to newly synthesized heme (**Fig. 1c**). By contrast, the HS1 eGFP/mKATE2 ratios in cells continuously maintained with or without SA remain characteristically high or low, respectively (**Fig. S2**).

The change in sensor heme saturation following the re-initiation of heme synthesis reveals three distinct phases: a lag phase, an exponential phase, and a stationary phase (**Fig. 1c**). The lag phase can be interpreted as the time required to alleviate the SA-mediated block in heme synthesis. The exponential phase represents the rate of heme binding to HS1, which is governed by the relative rates of heme synthesis and trafficking to the different subcellular locations. However, when comparing HS1 heme saturation kinetics between different compartments within a given strain, the data strictly represent the inter-compartmental heme trafficking rates since the contribution from heme synthesis is a constant. Indeed, expression of cytosolic, nuclear, or mitochondrial HS1 does not perturb heme synthesis (**Fig. S4a**). Since the final step of heme synthesis occurs with the insertion of ferrous iron into protoporphyrin IX, a reaction catalyzed by ferrochelatase at the matrix side of the mitochondrial IM, the heme trafficking rates reflect heme mobilization from the mitochondrial IM to the matrix, cytosol, or nucleus. Given the high affinity of HS1 for both ferric and ferrous heme, *K*_D_^III^= 10 nM and *K*_D_^II^ < 1 nM at pH 7.0^19^, and the negligible differences in FRET between eGFP and ferric and ferrous heme^19^, we cannot resolve the two oxidation states of heme. The stationary phase represents the maximum limiting heme saturation of the sensor 4 hours after alleviating the SA-mediated block in heme synthesis and typically spans ~70-100%. Due to the non-linearity in the fluorescence response of the sensor to heme above 80% bound and below 20% bound, we cannot easily resolve differences between sensor fractional saturation values that are > 80% and < 20%^1^.

The kinetics of heme binding to HS1 (**Fig. 1c**) can be fit to the logistic function described in **Equation 2**, where [A] is the maximal value of HS1 fractional saturation (amplitude), *k* is the first order rate constant (min^−1^), *t* is the time (min), and *t*_1/2_ is the midpoint of the transition. The lag time can be defined as *t*_1/2_ – 2/*k* ^22^.

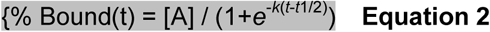

Quite surprisingly, the kinetics of HS1 heme saturation indicate that heme trafficking to locations as disparate as the mitochondrial matrix, cytosol, and nucleus are similar, with heme transport to the nucleus exhibiting a modest ~20% increase in rate constant relative to heme transport to the cytosol and mitochondrial matrix (**Fig. 1c**, **Table 1**). The remarkable similarity in the observed rates of heme trafficking to different locales might suggest that heme binding to the sensor is rate limiting. However, measurements of the rate of heme binding to HS1 in cell lysates of WT cells depleted of heme indicate that heme binding to the cytosolic, mitochondrial, and nuclear sensors occurs in less than 2 minutes (**Fig. S5**), which is much faster than the ~90-120 minutes it takes 50% of: (i) total cellular heme to be synthesized (**Fig. S4a**) or (ii) HS1 to become heme saturated in the SA pulse-chase assays (**Fig. 1c**, **Fig. S2**).

In order to rule out artifacts associated with heme sensor expression itself perturbing the observed heme trafficking rates, we expressed cytosolic HS1 under the control of weak (p*ADH1*), medium (p*TEF1*), or strong (p*GPD*) promoters and found that the observed heme trafficking rates were not affected despite a nearly 10-fold span in sensor expression (**Fig. S4b**). These results are generally consistent with our previous findings that sensor expression does not perturb heme homeostasis or otherwise affect cell viability^19^. Analogous experiments titrating expression with mitochondrial and nuclear sensors could not be completed due to low sensor expression and correspondingly low sensor signal to noise ratios associated with p*TEF1* or p*ADH1* driven expression.

In order to rule out artifacts associated with SA treatment, we re-capitulated the heme transport kinetics results by monitoring inter-compartmental heme trafficking kinetics in *hem1*∆ cells pulsed with a bolus of ALA, the product of the reaction catalyzed by ALA synthase (Hem1), to re-initiate heme synthesis. As shown in **Fig. 1d**, heme trafficking kinetics to the matrix, cytosol, and nucleus in *hem1*∆ cells are qualitatively similar to the results obtained from the SA pulse-chase assay using WT cells (**Fig. 1c**), with transport to the nucleus exhibiting a nearly 2-fold increase in rate constant relative to heme transport to the cytosol and mitochondrial matrix (**Table 1**). The absolute differences in heme transport kinetics between the SA pulse chase assay and feeding *hem1*∆ cells ALA likely reflects the differences associated with alleviating the SA block versus ALA uptake and assimilation into the heme biosynthetic pathway, respectively.

Given that the final step of heme synthesis occurs on the matrix side of the mitochondrial IM, our expectation was that the mitochondrial matrix would populate with heme first, followed by the cytosol and then the nucleus. However, the data indicate that once heme is synthesized in the mitochondrial IM, it disperses to multiple compartments almost simultaneously, suggesting the existence of parallel routes for heme mobilization to distinct locales. Of particular significance is the existence of a direct route for mitochondrial-nuclear heme trafficking given that the rates of heme trafficking from the mitochondrial IM to the nucleus are faster than to the mitochondrial matrix and cytosol (**Fig. 1c** and **1d**).

### Mgm1 and Dnm1 are positive and negative regulators of mitochondrial-nuclear heme trafficking, respectively

Our data indicate that heme is trafficked to the nucleus at faster rates than to the cytosol or even the mitochondrial matrix. What is the mechanism for mitochondrial-nuclear heme shuttling? The mitochondrial network is highly dynamic and is constantly remodeled by fusion and fission events. The dynamic behavior of the mitochondrial network is thought to be responsible for the proper cellular distribution and trafficking of a number of mitochondrial-derived metabolites, including various lipids^23,24^. For this reason, we hypothesized that mitochondrial dynamics may enable the facile distribution of heme, a lipid-like mitochondrial-derived metabolite^2^, to other organelles, including the nucleus^25,26^. To test this hypothesis, we perturbed genes involved in mitochondrial fusion and fission and assessed their impact on inter-compartmental heme trafficking kinetics using the SA-based pulse-chase assay described above.

#### Mitochondrial Fusion

Mitochondrial fusion occurs at both the IM and OM in coordinated but physically separable steps. In yeast, a pair of dynamin-like GTPases, Fzo1 and Mgm1, drive OM and IM fusion, respectively ^27^. In order to coordinate double membrane fusion, a protein spanning the mitochondrial intermembrane space, Ugo1, physically tethers Fzo1 and Mgm1, albeit the mechanisms orchestrating OM and IM fusion are not fully understood. Mgm1 exists in equilibrium between long (l-Mgm1) and short (s-Mgm1) isoforms, which are generated upon proteolytic cleavage by the rhomboid protease Pcp1^28,29^. It is thought that l-Mgm1, which has an inactive GTPase domain, acts as an anchor in the IM and interacts with and activates the GTPase activity of s-Mgm1 in the intermembrane space to promote IM fusion^28,29^. In mammals, there are homologs of Fzo1 (the mitofusins^30^), Mgm1 (OPA1^31^), and Ugo1 (SLC25A46^32^). Of note, in mammals, proteolytic processing of OPA1 requires action of the proteases YME1L and OMA1^33^.

We first tested the role of Mgm1 in regulating inter-compartmental heme trafficking. Relative to WT cells, the *mgm1*∆ strain exhibits a marked defect in nuclear heme trafficking, with a ~50% reduction in HS1 heme loading (**Fig. 2a**, **Table 1**). On the other hand, heme trafficking to the mitochondrial matrix and cytosol is not significantly impacted in *mgm1*∆ cells. Interestingly, heme availability to HS1 in the nucleus is similar between WT and *mgm1*∆ cells after prolonged growth, ~16 hours, suggesting that there are compensatory mechanisms that overcome the defects in nuclear heme trafficking in *mgm1*∆ cells (**Fig. S6a**). The rate of heme synthesis is un-perturbed by loss of Mgm1, indicating that the nuclear trafficking defect is not related to general defects in heme synthesis (**Fig. S6b**). The effects of the *mgm1*∆ mutant on nuclear heme trafficking is not due to the loss of mitochondrial DNA, which occurs with high frequency in *mgm1*∆ cells; *rho*^0^ cells that lack mitochondrial DNA do not exhibit an appreciable defect in nuclear heme trafficking (**Fig. 2b**, **Table 1**).

**Figure 2.**
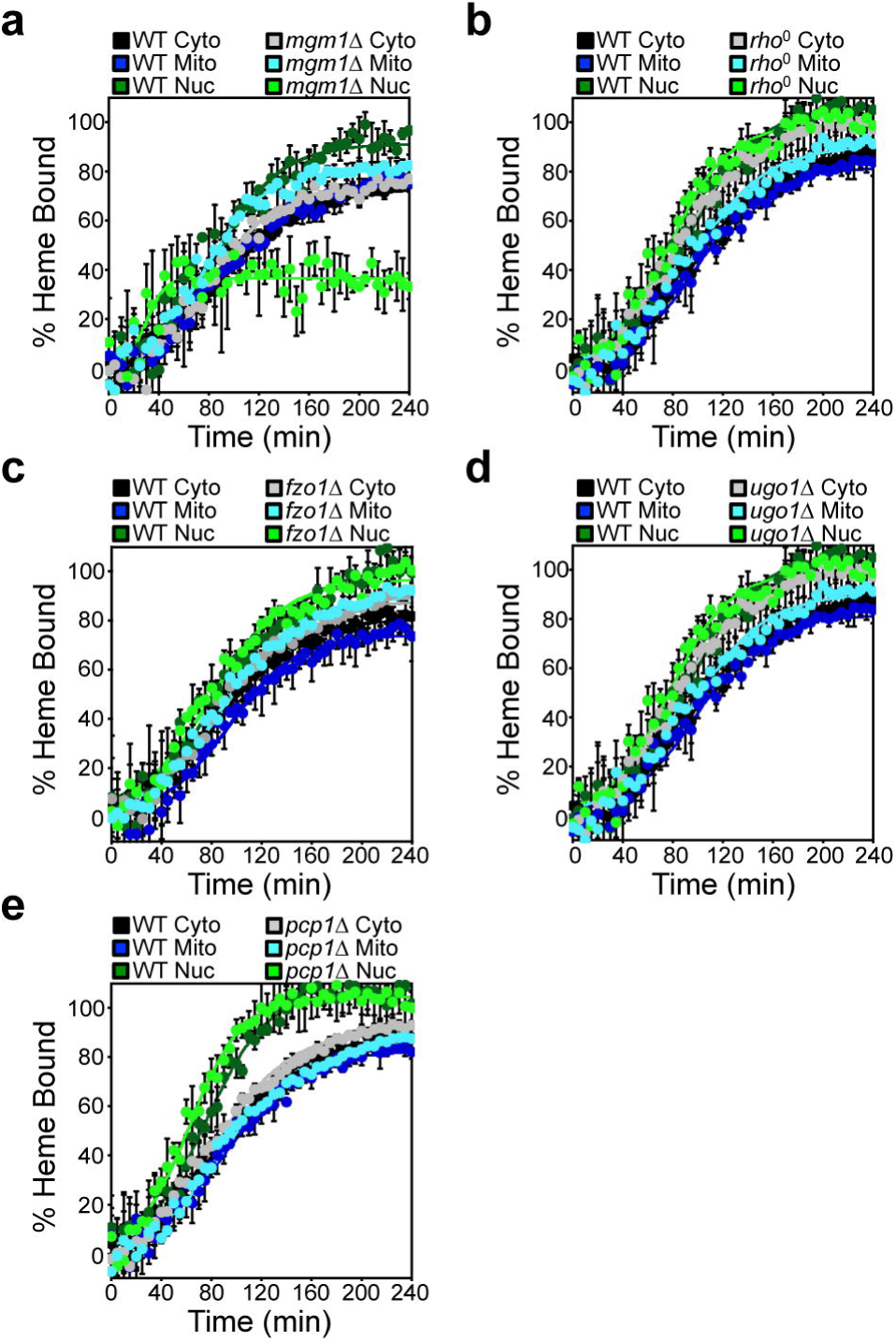
Mgm1 is a positive regulator of mitochondrial-nuclear heme trafficking. Inter-compartmental heme trafficking rates were monitored as in **Fig. 1c** using the SA pulse-chase assay for (**a**) *mgm1*∆, (**b**) *rho*^0^, (**c**) *fzo1*∆, (**d**) *ugo1*∆, and (**e**) *pcp1*∆ cells. The data represent the mean ± SD of independent triplicate cultures and the kinetic parameters derived from the fits to the data using equation 1 are indicated in **Table 1**.

In order to determine if the nuclear heme trafficking defect observed in *mgm1*∆ cells is specific to Mgm1 or more generally due to ablation of mitochondrial fusion, we tested other essential components of the mitochondrial fusion machinery, including Fzo1 and Ugo1. Interestingly, the *fzo1*∆ (**Fig. 2c**, **Table 1**) and *ugo1*∆ (**Fig. 2d**, **Table 1**) mutants do not exhibit defects in mitochondrial-nuclear heme trafficking, indicating that general perturbations to mitochondrial fusion do not impact nuclear heme trafficking.

Since the regulation of nuclear heme transport is specific to Mgm1, we tested a mutant with altered Mgm1 function. *pcp1*∆ cells are defective in fusion due to an inability to proteolytically convert l-Mgm1 to its short isoform. As seen in **Fig. 2e**, *pcp1*∆ cells do not exhibit a defect in mitochondrial-nuclear heme trafficking (**Table 1**). These data indicate that full-length Mgm1 regulates mitochondrial-nuclear heme transport, but the short Mgm1 isoform is largely dispensable. Altogether, our data strongly suggest that Mgm1, but not mitochondrial fusion *per se*, positively regulates mitochondrial-nuclear heme trafficking (**Fig. 2** and **Table 1**).

#### Mitochondrial Fission

In yeast, mitochondrial fission involves recruitment of the GTPase Dnm1 to its receptor Fis1 on the mitochondrial outer-membrane (OM), which is dependent on paralogous adapter proteins Mdv1 and Caf4^27^. Once assembled and oligomerized on the OM, Dnm1-catalyzed GTP hydrolysis drives the constriction and scission of mitochondrial tubules. In mammals, the Dnm1 homolog DRP1 drives fission, however the adapter proteins are less defined and there appear to be other mitochondrial receptors in addition to Fis1 ^27^.

We first tested the role of Dnm1 in regulating inter-compartmental heme trafficking. Most interestingly, relative to WT cells, the *dnm1*∆ strain exhibits an increase in the rate of heme trafficking to the mitochondrial matrix, cytosol, and nucleus, with the most pronounced effect on nuclear heme trafficking; .055 (.003) min^−1^ in *dnm1*∆ cells vs. .038 (.002) min^−1^ in WT cells (**Fig. 3a**, **Table 1**). After prolonged growth, 16 hours, heme availability to HS1 in all three compartments is similar between WT and *dnm1*∆ cells (**Fig. S6a**). Additionally, the rate of heme synthesis is unperturbed by loss of Dnm1, indicating that the increase in the rate of heme trafficking is not due to an increase in the rate of heme synthesis (**Fig. S6c**).

**Figure 3.**
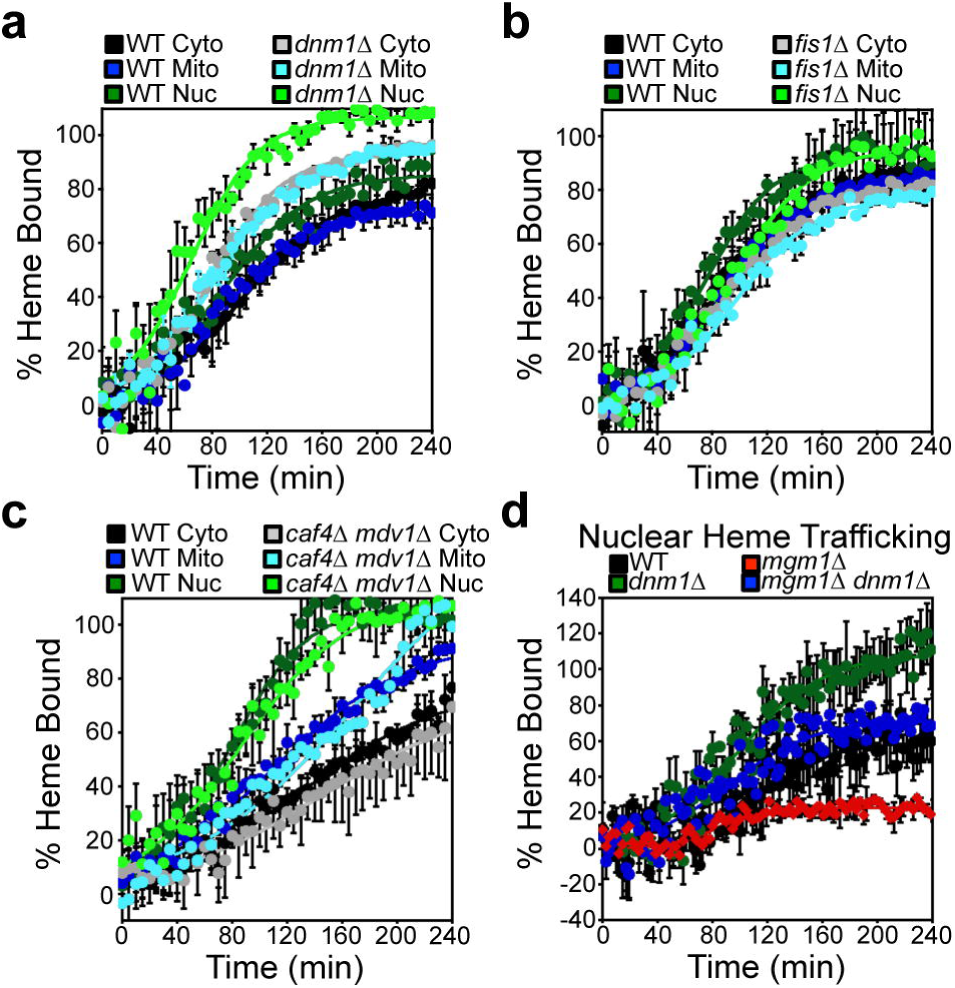
Dnm1 is a negative regulator of mitochondrial-nuclear heme trafficking. Inter-compartmental heme trafficking rates were monitored as in **Fig. 1c** using the SA pulse-chase assay for (**a**) *dnm1*∆, (**b**) *fis1*∆, and (**c**) *caf4*∆ *mdv1*∆ cells, and (**e**) *mgm1*∆ *dnm1*∆ cells. Fluorimetry data represent the mean ± SD of three independent cultures and the kinetic parameters derived from the fits to the data using equation 1 are indicated in **Table 1**.

In order to determine if the increased rate of nuclear heme trafficking observed in *dnm1*∆ cells is specific to Dnm1 or more generally due to ablation of mitochondrial fission, we tested other essential components of the mitochondrial fission machinery, including Fis1, Caf4, and Mdv1. Interestingly, *fis1*∆ (**Fig. 3b**, **Table 1**) and *caf4*∆ *mdv1*∆ (**Fig. 3c**, **Table 1**) mutants do not exhibit perturbations to mitochondrial-nuclear heme trafficking. Altogether, our data indicate that Dnm1, but not mitochondrial fission per se, negatively regulates mitochondrial-nuclear heme trafficking (**Fig. 3** and **Table 1**).

We next sought to determine the consequence of ablating both Mgm1 and Dnm1 on mitochondrial-nuclear heme trafficking. Most interestingly, *mgm1*∆ *dnm1*∆ double mutants exhibit WT-like rates of mitochondrial-nuclear heme trafficking (**Fig. 3d**). Therefore, in a manner analogous to the regulation of mitochondrial fusion and fission by Mgm1 and Dnm1, respectively, Mgm1 and Dnm1 regulate heme trafficking in opposing directions with similar magnitude.

### Mgm1 regulates the activation of the nuclear heme-regulated transcription factor Hap1

Given that Mgm1 and Dnm1 are positive and negative regulators of mitochondrial-nuclear heme trafficking, we sought to determine their impact on the activation of the heme-regulated transcription factor Hap1. Heme binding to Hap1 alters its ability to promote or repress transcription of a number of target genes, including *CYC1*, which Hap1 positively regulates^5,19,34-36^. In order to probe Hap1 activity, we used a transcriptional reporter that employs the promoter of a Hap1 target gene, p*CYC1*, driving the expression of enhanced green fluorescent protein (eGFP)^19^. Cells were first pre-cultured with 500 μM SA for 16 hours and, following this initial growth period, the SA conditioned cells were diluted into fresh media with (+) or without (∆) 500 μM SA for an additional 4 hours. A parallel set of cells continuously grown without SA (-) was cultured to provide a read out of steady-state Hap1 activity. As demonstrated in **Fig. 4**, not only is steady-state Hap1 activity greatly diminished in a heme deficient *hem1*∆ mutant and with SA conditioning (+), as expected, loss of Mgm1 results in a nearly 3-fold reduction in Hap1 activity. On the other hand, loss of Dnm1 does not affect basal Hap1 activity (**Fig. 4**, -). Since basal Hap1 activity in WT cells may already reflect heme saturation of Hap1, it is possible that the effects of the loss of Dnm1 are masked. To address this, we also measured Hap1 activity under non-heme saturating conditions in which Hap1 activity was probed just 4 hours after alleviation of the SA mediated block in heme synthesis. Under these conditions, *dnm1*∆ cells have increased Hap1 activity and *mgm1*∆ cells exhibit diminished Hap1 activity relative to WT cells (**Fig. 4**, ∆). Altogether, the positive and negative effects of Mgm1 and Dnm1 on nuclear heme trafficking, respectively, correlate with their effects on heme activation of Hap1.

**Figure 4.**
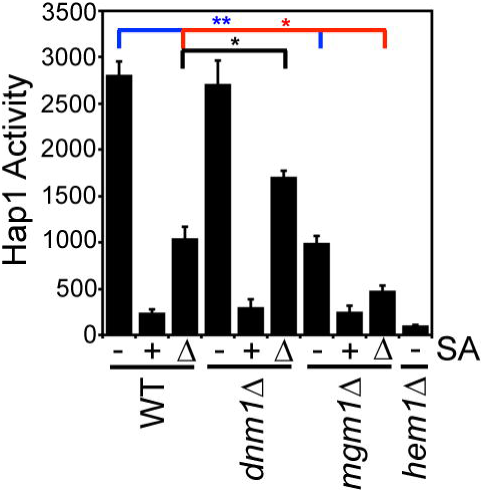
Mgm1 and Dnm1 regulate the activation of Hap1. Hap1p activity in the indicated strains as measured by a transcriptional reporter that used eGFP driven by the *CYC1* promoter, a Hap1p target gene. We report HAP1 activity in cells that were untreated with succinylacetone (SA) (-), treated with 500 μM SA (+), or were treated with 500 μM SA followed by shifting to media lacking SA for 4 hours (∆). Fluorimetry data represent the mean ± SD of three biological replicates; *p<0.05, **p<0.000005 by Student’s *t*-test.

### Heme regulates mitochondrial network morphology

Our results indicate that Mgm1 and Dnm1, factors that regulate mitochondrial dynamics, control mitochondrial-nuclear heme trafficking and heme activation of a heme-regulated transcription factor. We next asked if cellular heme levels could feedback to regulate mitochondrial network morphology and dynamics. Towards this end, by inhibiting heme synthesis using a 500 μM dose of SA for 4-6 hours, we found that there is a nearly 2-fold increase in mitochondrial fragmentation in SA-treated cells relative to control as early as 4 hours post-treatment (**Fig. 5a and 5b**). The number of cells with fragmented mitochondria further increases upon prolonged incubation with SA. The 500 μM dose of SA results in a ~60-80% decrease in intracellular heme (**Fig. 5c**) and alterations in mitochondrial network precede any appreciable changes in mitochondrial membrane potential (**Fig. 5d**). Hence, the SA-dependent increase in mitochondrial fragmentation is consistent with a role for heme in positively influencing mitochondrial fusion and/or negatively regulating mitochondrial fission. Future studies are warranted to investigate the molecular basis of this effect.

**Figure 5.**
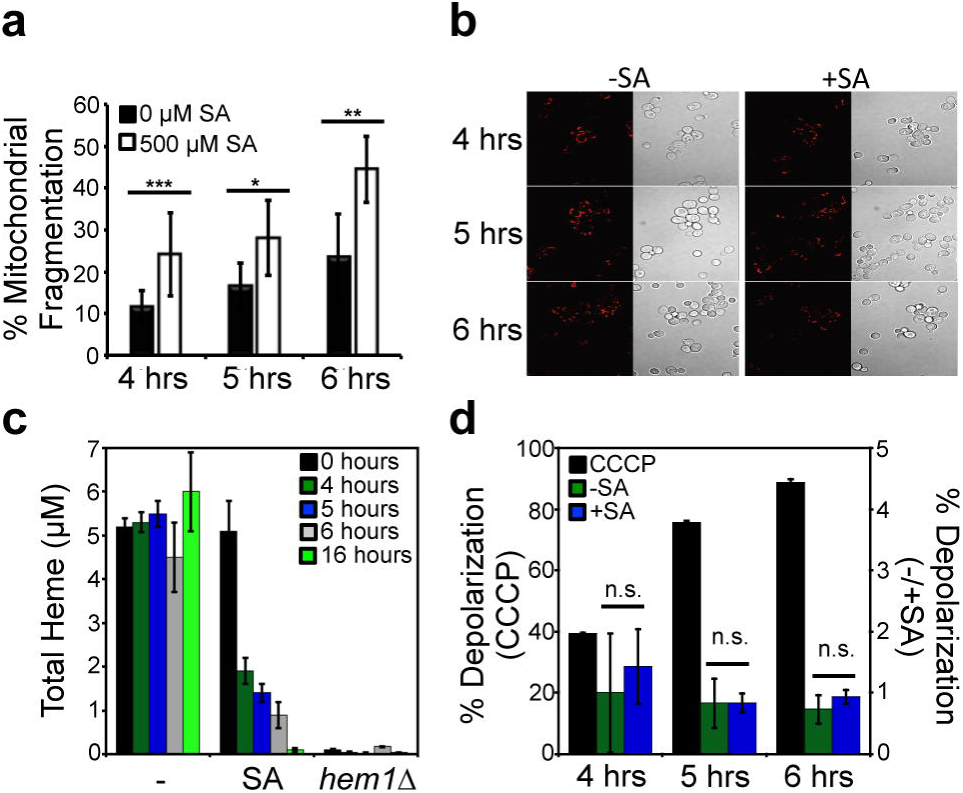
Heme regulates mitochondrial morphology. (**a**) Quantitative analysis of the mitochondrial network in Su9-RFP expressing wild type cells incubated with or without 500 μM succinylacetone (SA) for the indicated periods of time. Mitochondrial fragmentation was quantified as the ratio of the cells displaying punctate or dot-like morphology versus normal tubular mitochondrial networks. Data show mean values ± SD (n=3, with 300-400 cells per biological replicate); *p<0.05, **p<0.01, ***p<0.001 by Student’s *t*-test. (**b**) Representative images of the mitochondrial network from cells used to generate the data in panel **a**. (**c**) Cellular heme levels in wild type cells treated or not with 500 μM SA for an indicated period of time. Heme levels in the *hem1*∆ mutant served as a negative control. Data represent the mean ± SD of three biological replicates. (**d**) Changes in mitochondrial membrane potential in wild type cells treated with 500 μM SA (+SA) or 45 μM uncoupler CCCP (CCCP), or left untreated (-SA) for the indicated periods of time. Cells were stained with 200 nM membrane potential-sensitive dye DiOC_6_ and analyzed by flow cytometry using the FITC channel. Data show the mean values ± SD of 3 independent experiments; n.s., not significant by *t-*test.

## Discussion

The trafficking of heme from its site of synthesis to hemoproteins present in virtually every subcellular compartment underlies all heme-dependent processes. However, the molecules and mechanisms that mediate heme mobilization, as well as its spatio-temporal dynamics, remain poorly understood^1,2^. Herein, we developed a new quantitative pulse-chase assay in which, upon the initiation of heme synthesis, genetically encoded heme sensors were used to monitor the flow of heme from the matrix-side of the mitochondrial IM, where the final step of heme synthesis occurs, to the mitochondrial matrix, cytosol, and nucleus. Most surprisingly, we found that mitochondrial-nuclear heme trafficking occurs at faster rates than to the matrix or cytosol, which acquire heme at equal rates (**Fig. 1c** and **1d**). These results suggest the existence of multiple parallel pathways for heme mobilization to different cellular locales; thereby challenging the existing paradigm and biochemical intuition that heme sequentially populates various subcellular locations. Moreover, using molecular genetic approaches, we found that GTPases involved in mitochondrial fusion, Mgm1, and fission, Dnm1, are positive and negative regulators of mitochondrial-nuclear heme trafficking, respectively (**Fig. 2a** and **3a**).

Our results indicating that there is a direct pathway for mitochondrial-nuclear heme shuttling can be considered in the context of four emerging paradigms for conceptualizing the utilization of mitochondrial-derived heme^2^, namely: **1)** heme is transported into the cytosol via heme exporter proteins where it contributes to a kinetically labile pool that can be mobilized to hemoproteins throughout the cell; **2)** heme trafficking factors physically associate with the heme biosynthetic machinery in order to facilitate the distribution of heme, **3)** mitochondrial dynamics and contact sites with other organelles facilitates the exchange and distribution of heme between organelles; and/or **4)** mitochondrial-derived vesicles encapsulate and traffic heme^2^. While significant experimental evidence exists for mechanisms “1” and “2”, less data is available to support mechanisms “3” and “4”.

With respect to mechanism “1”, mammalian Flvcr1b, an isoform of feline leukemia virus subgroup C cellular receptor 1, was recently proposed to be a mitochondrial heme exporter and was required for erythropoiesis and hemoglobinization^37^. Once heme is outside the mitochondria, heme trafficking factors may then ferry heme to different cellular locations to populate *apo*-hemoproteins. For example, GAPDH has been found to not only buffer cytosolic labile heme^19^, but also regulate heme trafficking to hemoproteins like nitric oxide synthase^38^ and heme-regulated transcription factor Hap1^19^. However, since Flvcr1 is not conserved in lower eukaryotes like yeast^2^, there are likely other conserved pathways for mitochondrial heme efflux. Our data suggest that mitochondrial-derived heme is not distributed sequentially, but rather through parallel pathways of distribution. In particular, we identified a direct route for mitochondrial-nuclear heme trafficking that is Mgm1-dependent.

With respect to mechanism “2”, recent studies have demonstrated that the heme biosynthetic machinery exists as a multi-protein supracomplex or metabolon, and putative heme trafficking factors PGRMC1/2 were found to physically associate with the heme metabolon^39,40^. These findings suggest that heme trafficking factors can directly accept heme from the terminal enzyme in the heme biosynthetic pathway, ferrochelatase, potentially bypassing the labile heme pool, prior to delivering heme to cognate hemoproteins. Given that the human homolog of Mgm1, OPA1, was previously found to physically associate with ferrochelatase in the heme metabolon^40^, it is tempting to speculate that Mgm1/OPA1 may be also accepting heme from ferrochelatase prior to delivery to the nucleus. Our efforts are now focused on elucidating the molecular mechanisms underlying Mgm1-mediated nuclear heme trafficking.

Scenario “3” and “4”-based mechanisms are motivated by studies on lipid trafficking, albeit there is scant experimental evidence for their respective roles in trafficking heme, a lipid-like metabolite^2^. Notably, the putative heme trafficking factor PGRMC1 has recently been identified as part of the mammalian ER-mitochondrial tethering complex^41^, thus direct interactions between mitochondria and ER may facilitate heme trafficking through out the cell, including to the nucleus. Since mitochondrial fission and fusion factors have been physically and functionally linked to mitochondrial-ER contact sites^25^, we speculate that Dnm1/DRP1 and Mgm1/OPA1 may also act to regulate heme trafficking via the ER.

The dynamic mitochondrial network, which is constantly being re-shaped by fission and fusion events, is thought to be responsible for the proper distribution and trafficking of mitochondrial-derived metabolites, including certain lipids^23,24^. However, we find that dynamic mitochondria itself is not a requirement for normal mitochondrial-nuclear heme trafficking. Indeed, an *mgm1*∆ *dnm1*∆ mutant that cannot undergo fission and fusion, but that exhibits normal mitochondrial morphology^42^ (**Fig. S7**), exhibits WT-like rates of heme trafficking to the nucleus (**Fig. 3d**).

How might Dnm1 act as a negative regulator of heme trafficking? This GTPase exists in an equilibrium between membrane-bound and soluble oligomeric species^43^. Binding of lipids such as cardiolipin, can stimulate GTPase activity and Dnm1/Drp1-dependent membrane fission events^43^. Given heme’s lipid-like properties^2^, we speculate that heme may be sequestered by certain Dnm1 species to suppress heme trafficking.

In such a model, Mgm1 and Dnm1 would cooperate to regulate the appropriate flux of heme to the nucleus. We are currently exploring the molecular mechanisms underlying Dnm1-mediated negative regulation of heme trafficking and how it may cooperate with Mgm1 to control heme flux to the nucleus.

Our results also have significant implications for the mechanisms underlying heme-based mitochondrial-nuclear retrograde signaling^44^. Given that heme is a mitochondrial-derived metabolite and that there are heme-regulated transcription factors present in all eukaryotes, it has long been posited that heme could act as a retrograde signal that can communicate mitochondrial metabolic status to the nucleus^1,2^. Our results indicate that there is a direct pathway for mitochondrial-nuclear heme trafficking to facilitate the activation of heme-regulated transcription factors.

A number of human diseases, including certain cancers, neurodegenerative disorders, and blood diseases, are associated with both defects in heme homeostasis and mitochondrial fragmentation, which can occur as a result of altered fission or fusion^45,46^. Our results indicating GTPases that regulate mitochondrial dynamics also regulate heme trafficking and heme itself is required to prevent mitochondrial fragmentation suggest that heme homeostasis and mitochondrial dynamics are integrally linked and may have disease implications. For instance, as one example, in Alzheimer’s disease, amyloid beta (Aβ) over-expression results in increased mitochondrial fragmentation^47^. Given previous studies indicating that Aβ can bind and sequester cellular heme^15,16^, our finding that heme suppresses mitochondrial fragmentation suggests that Aβ may in part promote mitochondrial fragmentation by limiting heme availability.

Altogether, our *in vivo* approach to monitor real-time dynamics of inter-compartmental heme trafficking coupled with molecular genetic approaches have uncovered fundamental aspects of the mechanisms underlying heme mobilization and utilization.

## Materials and Methods

All yeast strains, culture conditions, reagents, and experimental protocols are described in the **Supporting Information**.

## Acknowledgements

This work was supported by the National Institutes of Health (ES025661 to ARR, GM108975 to OK, DK111653 to AEM), the National Science Foundation (MCB-1552791 to ARR), the Blanchard Professorship (to ARR), Georgia Institute of Technology (to ARR), the U.S. Department of Education GAANN Program (Grant P200A120081 to OMG), and the University of Nebraska Molecular Mechanisms of Disease graduate program (to JVD). We wish to acknowledge the core facilities at the Parker H. Petit Institute for Bioengineering and Bioscience at the Georgia Institute of Technology for use of the shared equipment, services, and expertise.

Author Contributions
OMG, OK, AEM, and ARR designed research. OMG, JVD, IB, and ARR performed research. All authors analyzed and interpreted data. OMG and ARR wrote the paper with input from all authors.

