## Supporting Information for "Mitochondrial-nuclear heme trafficking is regulated by GTPases that control mitochondrial dynamics"

EXPERIMENTAL MODEL

- Yeast Strains, Transformations, and Growth Conditions
- Plasmids

METHOD DETAILS

- Heme Trafficking Dynamics Assay
- Total Heme Quantification
- Hap1 Activity
- Isolation of Mitochondria and Nuclei to Confirm Sensor Localization
- Confirmation of Heme Sensor Localization by Microscopy
- Confirmation of Mitochondrial Morphology in Fission and Fusion Mutants by Microscopy
- Mitochondrial Fragmentation Assay
- Mitochondrial Membrane Potential Assay

SUPPLEMENTAL FIGURE LEGENDS

SUPPLEMENTAL REFERENCES

**EXPERIMENTAL MODEL**

**Yeast Strains, Transformations, and Growth Conditions**

*S. cerevisiae* strains used in this study were derived from BY4741 (MATa, *his3*Δ1, *leu2*Δ0, *met15*Δ0, *ura3*Δ0). *fis1*∆::*Kan*MX4, *dnm1*∆::*Kan*MX4, *mgm1*∆::*Kan*MX4, *pcp1*∆::*Kan*MX4, *ugo1*∆::*Kan*MX4, *caf4*∆*::KanMX4*, *mdv1*∆*::KanMX4* strains were obtained from the yeast gene deletion collection (Thermo Fisher Scientific). We also utilized the previously reported strains, LJ109 (rho^0^)^1^ and DH001b-3 (*hem1*∆::*HIS3*)^2^. AR1029-3 (*mdv1*Δ::*Kan*MX4 *caf4*Δ::*HIS3*) was generated by deleting *CAF4* with pAR1047 in *mdv1*Δ::*Kan*MX4 cells. OM232 (*mgm1*Δ::*HIS3*) and OM233 (*dnm1*Δ::*Kan*MX4 *mgm1*Δ::*HIS3*) was generated by deleting *MGM1* with pAR1051 in WT and *dnm1*∆*::KanMX4* cells, respectively. All strains were confirmed by PCR, mitochondrial morphology (**Fig. S7**), and, if derived from the yeast deletion collection, sequencing the unique barcodes up and downstream of the *KanMX4* deletion cassette.

Yeast transformations were performed by the lithium acetate procedure ^3^. Strains were maintained at 30° C on either enriched yeast extract (1%)-peptone (2%) based medium supplemented with 2% glucose (YPD), or synthetic complete medium (SC) supplemented with 2% glucose and the appropriate amino acids to maintain selection ^2^. Cells cultured on solid media plates were done so with YPD or SC media supplemented with 2% agar ^2^. Selection for yeast strains containing the KanMX4 marker was done with YPD agar plates supplemented with G418 (200 μg/mL) ^2^. WT cells treated with the heme synthesis inhibitor, succinylacetone (SA), and *hem1*Δ cells were cultured in YPD or SC media supplemented with 50 μg/mL of 5-aminolevulinic acid (ALA) or 15 mg/mL of ergosterol and 0.5% Tween-80 (YPDE or SCE, respectively)^2,4^. All liquid cultures were maintained at 30 °C and shaken at 220 RPM.

**Plasmids**

The *caf4*::*HIS3* disruption plasmid, pAR1047, was generated by first PCR amplifying the upstream (-650 to -129) and downstream (+2110 to +2479) sequences relative to the *CAF4* translational start site, introducing 5’ BamHI / 3’ XhoI and 5’ XbaI / 3’ BamHI restriction sites, respectively. The *CAF4* PCR products were digested with the enzymes indicated and ligated in a trimolecular reaction into the *HIS3* integrating plasmid pRS403^5^ digested with XhoI and XbaI, resulting in pAR1047. Transformation of yeast strains with pAR1047 linearized with BamHI resulted in deletion of *CAF4* sequences from -128 to +2109.

The *mgm1*::*HIS3* disruption plasmid, pAR1051, was generated by first PCR amplifying the upstream (-532 to -5) and downstream (+2736 to +3023) sequences relative to the *MGM1* translational start site, introducing 5’ BamHI / 3’ XhoI and 5’ XbaI / 3’ BamHI restriction sites, respectively. The *MGM1* PCR products were digested with the enzymes indicated and ligated in a trimolecular reaction into the *HIS3* integrating plasmid pRS403^5^ digested with XhoI and XbaI, resulting in pAR1051. Transformation of yeast strains with pAR1051 linearized with BamHI resulted in deletion of *MGM1* sequences from -4 to +2735.

Cytosolic, mitochondrial, and nuclear-targeted heme sensors, HS1, were sub-cloned into pRS415 and driven by *ADH*, *TEF*, or *GPD* promoters as previously described^2^. The Hap1 reporter plasmid in which eGFP is driven by the *CYC1* promoter was also previously described^2^.

**METHOD DETAILS**

**Heme Trafficking Dynamics Assay**

*Overview.* Inter-compartmental heme trafficking rates were monitored by: **a.** inhibiting heme synthesis with succinylacetone (SA) in sensor expressing cells; **b.** removing the block in heme synthesis by re-suspending cells into media lacking SA; and **c.** monitoring the time-dependent change in the percentage of heme bound to heme sensor 1 (HS1) localized to the cytosol, nucleus, and mitochondrial matrix upon the re-initiation of heme synthesis. The percentage of heme bound to the heme sensor (% Bound) is calculated using Equation S1^2^; where *R* is the HS1 eGFP/mKATE2 ratio and *R*_max_ and *R*_min_ is the eGFP/mKATE2 fluorescence ratio when the sensor is 100% or 0% bound to heme, respectively.

[% Bound] = ([*R* – *R*_min_] / [*R*_max_ – *R*_min_])*100	**Equation S1**

*R*_min_ is determined by measuring the HS1 eGFP/mKATE2 ratio in parallel cultures that are conditioned with succinylacetone (SA), which inhibits the second enzyme in the heme biosynthetic pathway, 5-aminolevulinic acid (ALA) dehydratase (ALAD)^6^, and *R*_max_ is determined by measuring the HS1 eGFP/mKATE2 ratio in parallel WT cultures grown without SA^2^. Background fluorescence from cells not expressing HS1 was subtracted from the eGFP (ex. 488 nm, em. 510 nm) and mKATE2 (ex. 588 nm, em. 620 nm) channels of sensor expressing cells.

*Growth for SA-pulse chase assay.* HS1-expressing cells were cultured with or without 500 μM SA (Sigma-Aldrich) in SCE-LEU media. Triplicate 5 mL cultures were seeded at an initial optical density of OD_600nm_ = .01-.02 (2-4 x 10^5^ cells/mL) and grown for 14-16 hours at 30 °C and shaking at 220 RPM until cells reached a final density of OD_600nm_ ~ 1.0 (2 x 10^7^ cells/mL). After culturing, 1 OD or 2 x 10^7^ cells were harvested, washed twice with 1 mL of ultrapure water, and resuspended in 1 mL of fresh SC-LEU media. The cells that were pre-cultured without SA provided HS1 *R*_max_ values. The SA-conditioned cells were split into two 500 μL fractions. One fraction was treated with 500 μM SA to give HS1 *R*_min_ values. The other fraction was not treated with SA so that heme synthesis could be re-initiated to give compartment-specific heme trafficking rates. HS1 fluorescence was monitored on 200 uL of a 1 OD/mL (2 x 10^7^ cells/mL) cell suspension using black Greiner Bio-one flat bottom fluorescence plates and a Synergy Mx multi-modal plate reader. eGFP (ex. 488 nm, em. 510 nm) and mKATE2 (ex. 588 nm, em. 620 nm) fluorescence was recorded every 5 minutes for 4 hours, with the plate being shaken at medium-strength for 30 seconds prior to each read. Background fluorescence of cells not expressing the heme sensors were recorded and subtracted from the eGFP and mKATE2 fluorescence values.

*Growth for ALA pulse-chase assay.* HS1-expressing *hem1*∆ cells were cultured with or without 400 μM 5-aminolevulinic acid (ALA) (Sigma-Aldrich) in SCE-LEU media. Triplicate 5 mL cultures were seeded at an initial optical density of OD_600nm_ = .01-.02 (2-4 x 10^5^ cells/mL) and grown for 14-16 hours at 30 °C and shaking at 220 RPM until cells reached a final density of OD_600nm_ ~ 1.0 (2 x 10^7^ cells/mL). After culturing, 1 OD or 2 x 10^7^ cells were harvested, washed twice with 1 mL of ultrapure water, and resuspended in 1 mL of fresh SC-LEU media. The cells that were pre-cultured with ALA provided HS1 *R*_max_ values. The cells that were cultured without ALA were split into two 500 μL fractions. One fraction was used to give HS1 *R*_min_ values. The other fraction was treated with 400 μM ALA so that heme synthesis could be initiated to give compartment-specific heme trafficking rates. HS1 fluorescence was monitored on 200 uL of a 1 OD/mL (2 x 10^7^ cells/mL) cell suspension using black Greiner Bio-one flat bottom fluorescence plates and a Synergy Mx multi-modal plate reader. eGFP (ex. 488 nm, em. 510 nm) and mKATE2 (ex. 588 nm, em. 620 nm) fluorescence was recorded every 5 minutes for 4 hours, with the plate being shaken at medium-strength for 30 seconds prior to each read. Background fluorescence of cells not expressing the heme sensors were recorded and subtracted from the eGFP and mKATE2 fluorescence values.

It should be noted that due to evaporation and drying of cell cultures in the plate reader, reliable measurements could not be achieved after 4 hours of continuous monitoring.

**Total Heme Quantification**

Measurements of total heme were accomplished using a fluorimetric assay designed to measure the fluorescence of protoporphyrin IX upon the release of iron from heme as previously described^7^. For all total heme measurements, ~2 x 10^8^ cells were harvested, washed in sterile ultrapure water, and resuspended in 500 μL of 20 mM oxalic acid and stored in a closed box at 4 °C overnight (16-18 hours). Next, an equal volume (500 μL) of 2 M oxalic acid was added to the cell suspensions in 20 mM oxalic acid. The samples were split, with half the cell suspension transferred to a heat block set at 95 °C and heated for 30 minutes and the other half of the cell suspension kept at room temperature (~25 °C) for 30 minutes. All suspensions were centrifuged for 2 minutes on a table-top microfuge at 21000 x g and the porphyrin fluorescence (ex: 400 nm, em: 620 nm) of 200 μL of each sample was recorded on a Synergy Mx multi-modal plate reader using black Greiner Bio-one flat bottom fluorescence plates. Heme concentrations were calculated from a standard curve prepared by diluting 500-1500 μM hemin chloride stock solutions in 0.1 M NaOH into ultrapure water, which was then added back to extra cell samples as prepared above. In order to calculate heme concentrations, the fluorescence of the unboiled sample (taken to be the background level of protoporphyrin IX) is subtracted from the fluorescence of the boiled sample (taken to be the free base porphyrin generated upon the release of heme iron). The cellular concentration of heme is determined by dividing the moles of heme determined in this fluorescence assay and dividing by the number of cells analyzed, giving moles of heme per cell, and then converting to a cellular concentration by dividing by the volume of a yeast cell, taken to be 50 fL^2^.

In order to measure heme synthesis rates, exponential phase cells were pre-conditioned with 500 μM SA for 14-16 hours in SCE-LEU media as described above for the heme trafficking dynamics assay. Following this, cells were washed and resuspended in media lacking SA and aliquots of cells were harvested and washed for total heme quantification as described above.

**Hap1 Activity**

Cells expressing p415-*CYC1*-*eGFP*, or *eGFP* driven by the Hap1 regulated *CYC1* promoter, were cultured in 50 mLs of SCE-LEU in 250 mL Erlenmeyer flasks, with or without 500 μM SA, for 14-16 hours to an optical density of OD_600nm_ ~ 1.0 (2 x 10^7^ cells/mL). Cells were washed with sterile ultrapure water and diluted into fresh SC-LEU media at an optical density of OD_600nm_ = 0.25 and allowed to grow for an additional 4 hours. The culture that was initially not conditioned with SA remained untreated during the 4-hour growth phase and these cultures represented basal Hap1 activity. The culture that was pre-conditioned with SA was split into two fractions. One fraction was treated with 500 μM SA for the 4-hour growth phase and these cultures served as a negative control, representing minimal Hap1 mediated activation of *CYC1*. The other fraction was not treated with SA for the 4-hour growth phase and these cultures represented Hap1 activity under conditions in which Hap1 was not saturated with heme. After growth for 4 hours, cells were washed in sterile ultrapure water and resuspended in PBS to a concentration of 1 x 10^8^ cells/mL and 100 uL was used to measure eGFP fluorescence (ex. 488 nm, em. 510 nm). Background auto-fluorescence of cells not expressing eGFP was recorded and subtracted from the p415-*CYC1*-*eGFP* expressing strains.

**Isolation of Mitochondria and Nuclei to Confirm Heme Sensor Localization**

Mitochondria and nuclei were isolated using yeast mitochondrial (Cat # K259-50) and nuclear (Cat # K289-50) isolation kits from BioVision according to the manufacturer’s instructions. All reagents were supplied by the isolation kits. For isolation of mitochondria, 50 mLs of sensor expressing cells were cultured in SC-LEU in 250 mL Erlenmeyer flasks to a final density of OD_600nm_ = 1.0 (2 x 10^7^ cells/mL). 4 x 10^8^ cells were harvested, washed in ultrapure water, and resuspended in “Buffer A” containing 10 mM DTT for 30 minutes at 30 °C with occasional gentle agitation. Cells were then harvested by centrifugation at 1,500 x g for 5 minutes at room temperature. Cells were then re-suspended in 1 mL of “Buffer B” containing manufacturer provided Zymolyase and incubated at 30 °C with occasional gentle inversion. Spheroplast formation was monitored by diluting 10 uL of the cell suspension into ultrapure water and monitoring the decrease in OD_600nm_ such that it was > 80% less than the initial value, which typically took 30-60 minutes. Spheroplasts were harvested by centrifugation at 1500 x g for 5 minutes, resuspended in “Homogenization Buffer” containing a protease inhibitor cocktail, and lysed with 15 strokes of a Dounce Homogenizer on ice. The lysate was centrifuged at 600 x g for 5 minutes at 4 °C to remove cell debris. The supernatant, which contained the mitochondria, was again centrifuged for 5 minutes at 600 x g to remove additional debris. Finally, mitochondria was harvested by centrifugation at 12000 x g for 10 minutes at 4 °C. The mitochondrial pellet was resuspended in 50 μL of “Storage Buffer”. In order to validate sensor mitochondrial localization, 5% of the total spheroplast fraction, mitochondrial fraction, and the post-mitochondrial fraction (supernatant from the 12000 x g centrifugation step) were electrophoresed on a 14% tris-glycine SDS-PAGE gel and immunoblotted using α-PGK1 (Life Technologies; Cat # PA528612; 1:5000 dilution), a cytosolic marker protein, α-porin (Life Technologies; Cat # 459500; 1:5000 dilution), a mitochondrial marker protein, and α-GFP (Genetex; Cat # GTX30738; 1:5000 dilution) to probe HS1 expression. A goat α-rabbit secondary antibody conjugated to a 680 nm emitting fluorophore (VWR/Biotium; Cat # 89138-520; 1:10000 dilution) was used to probe for PGK1 and GFP. A goat α-mouse secondary antibody conjugated to a 680 nm emitting fluorophore (VWR/Biotium; Cat # 89138-516; 1:10000 dilution) was used to probe for Porin. All gels were imaged on a LiCOR Odyssey Infrared imager^1,2^.

For isolation of nuclei, 50 mLs of sensor expressing cells were cultured in SC-LEU in 250 mL Erlenmeyer flasks to a final density of OD_600nm_ = 1.0 (2 x 10^7^ cells/mL). 4 x 10^8^ cells were harvested, washed in ultrapure water, and resuspended in “Buffer A” containing 10 mM DTT for 30 minutes at 30 °C with occasional gentle agitation. Cells were then harvested by centrifugation at 1,500 x g for 5 minutes at room temperature. Cells were then re-suspended in 1 mL of “Buffer B” containing manufacturer provided Zymolyase and incubated at 30 °C with occasional gentle inversion. Spheroplast formation was monitored by diluting 10 uL of the cell suspension into ultrapure water and monitoring the decrease in OD_600nm_ such that it was > 80% less than the initial value, which typically took 30-60 minutes. Spheroplasts were harvested by centrifugation at 1500 x g for 5 minutes, resuspended in 1 mL of “Buffer N” containing a protease inhibitor cocktail, and lysed with 5 strokes of a Dounce Homogenizer on ice. The suspension was incubated for 30 minutes at room temperature with gentle agitation every 3-5 minutes. The lysate was centrifuged at 1500 x g for 5 minutes at 4 °C to remove cell debris. Finally, nuclei were harvested by centrifugation at 20000 x g for 10 minutes at 4 °C. The nuclear pellet was resuspended in 100 μL of “Buffer N”. In order to validate sensor nuclear localization, 5% of the total spheroplast fraction, nuclear fraction, and the post-nuclear fraction (supernatant from the 20000 x g centrifugation step) were electrophoresed on a 14% tris-glycine SDS-PAGE gel and immunoblotted using α-PGK1 (Life Technologies; Cat # PA528612; 1:5000 dilution), a cytosolic marker protein, α-NOP1 (Life Technologies; Cat # MA110025; 1:10000 dilution), a nuclear marker protein, and α-GFP (Genetex; Cat # GTX30738; 1:5000 dilution) to probe HS1 expression. A goat α-rabbit secondary antibody conjugated to a 680 nm emitting fluorophore (VWR/Biotium; Cat # 89138-520; 1:10000 dilution) was used to probe for PGK1 and GFP. A goat α-mouse secondary antibody conjugated to a 680 nm emitting fluorophore (VWR/Biotium; Cat # 89138-516; 1:10000 dilution) was used to probe for NOP1. All gels were imaged on a LiCOR Odyssey Infrared imager^1,2^.

**Confirmation of Heme Sensor Localization by Microscopy**

To confirm mitochondrial or nuclear localization of the sensors, prior to microscopy, 1 x 10^7^ exponential phase cells were incubated with 2.0 μg/mL 4’, 6-diamidino- 2-phenylindole (DAPI) (Invitrogen) in SC-LEU media for 45-60 minutes to stain nuclear or mitochondrial DNA^2^. Laser scanning confocal microscopy was accomplished on a Zeiss ELYRA LSM 780 Super-resolution Microscope equipped with a 63x, 1.4 numerical aperture oil objective. DAPI was excited at 405 nm and emission was collected using 410-483 nm band pass filters. eGFP was excited with the 488 nm line of an argon ion laser, while mKATE2 was excited using the 594 nm of a HeNe laser line. The 491-588 nm and 599-690 nm band pass filters were used to filter emission for eGFP and mKATE, respectively. Images were collected using Zeiss software and analyzed with ImageJ 1.48v (Rasband, W.S., ImageJ, U. S. National Institutes of Health, Bethesda, Maryland, USA, http://rsb.info.nih.gov/ij/, 1997-2007). Auto-fluorescence of unlabeled cells was subtracted from the DAPI, eGFP, and mKATE2 channels to produce the finalized images in **Figure 1b**.

**Confirmation of Mitochondrial Morphology in Fission and Fusion**

To visualize the mitochondrial network, 1 x 10^7^ exponential phase cells were incubated with 500 μM Mito Tracker Red CM-H_2_XRos (Thermo Fisher) in SC-LEU media for 45-60 minutes. Confocal microscopy was done on a Zeiss ELYRA LSM 780 Super-resolution Microscope equipped with a 63x, 1.4 numerical aperture oil objective. Mito Tracker Red was excited using the 594 nm of a HeNe laser line and emission was collected using a 415-735 nm band pass filter. Images were collected using Zeiss software and analyzed with ImageJ 1.48v (Rasband, W.S.,ImageJ, U. S. National Institutes of Health, Bethesda, Maryland, USA, http://rsb.info.nih.gov/ij/, 1997-2007). The various mitochondrial network morphology types observed are outlined in **Fig. S7a** and ~50 cells from each strain were analyzed to confirm the expected mitochondrial morphology (**Fig. S7b**). A representative image of the mitochondrial network from each strain is depicted in **Fig. S7c**. Fission and fusion mutants have characteristic elongated or punctate mitochondrial networks, respectively.

**Mitochondrial Fragmentation Assay**

Mitochondrial morphology was assessed in wild-type yeast (W303, *MAT ade2-1 can1-100 his3-11,15 leu2-3,112 trp1-1 ura3-1*) transformed with the pYX142-Su9-RFP plasmid^8^. The pYX142-Su9-RFP plasmid expresses a mitochondria-targeted red fluorescent protein (RFP), allowing for visualization of mitochondria using a fluorescence microscope. Cells were grown overnight in synthetic complete (SC) medium supplemented with appropriate nutrients and 2% glucose as the sole carbon source. The cells were then sub-cultured into fresh pre-warmed medium at an OD_600_ of 0.2 for three hours, and then treated with succinylacetone (SA) at a final concentration of 500 μM and cultured for 4, 5, or 6 hours post-SA treatment. Imaging was performed using the Olympus IX81-FV5000 confocal laser-scanning microscope at 543-nm laser line with a 100x oil objective. Images were acquired and processed with Fluoview 500 software (Olympus America). Mitochondrial fragmentation was quantified based on the ratio of the cells displaying punctated or dotted-like morphology versus normal ribbon-like mitochondrial networks. At least 300 cells were analyzed for each sample per replicate.

**Mitochondrial Membrane Potential Assay**

The mitochondrial membrane potential was assayed on cells cultured in a similar manner to that described above for the assessment of heme-dependent changes in mitochondrial fragmentation using a BD FACSCanto^TM^ II system (Becton Dickinson) as previously described^9^. After culturing for the indicated times, cells were first washed with phosphate-buffered saline (PBS), and then stained with 200 nM 3,3′-dihexyloxacarbocyanine iodide (DiOC_6_) (Molecular Probes) for 30 min at 30 ^°^C. Finally, the cells were washed twice with PBS and analyzed by flow cytometry using the FL3 channel without compensation. The data were collected, analyzed and plotted using BD FACSDiva software v6.1.1 (Becton Dickinson).

**Supplemental Figure Legends**

**Fig. S1.** Confirmation of heme sensor, HS1, localization as assessed by cell fractionation and immunoblotting. Nuclei (**a**) and mitochondria (**b**) were isolated as described in the **Method Details**, and expression of HS1 was probed using α-GFP antibodies. The indicated fractions were confirmed by probing the expression of PGK1, NOP1, and POR1, which are cytosolic, nuclear, and mitochondrial marker proteins, respectively. 5% of the whole cell extract (WCE), derived from the spheroplast fraction, nuclear fraction (NF), mitochondrial fraction (MF), post-mitochondrial fraction (PMF), or post-nuclear fraction (PNF) were electrophoresed on a 14% tris-glycine SDS-PAGE gel.

**Fig. S2.** Representative “raw” kinetic traces of the heme trafficking dynamics assay. WT cells expressing HS1 in the cytosol (**a**), nucleus (**b**), or mitochondria (**c**) were depleted of heme using 500 μM succinylacetone (SA), a heme biosynthetic inhibitor, and, upon the re-initiation of heme synthesis, the rates of heme trafficking to the indicated subcellular locations were monitored by measuring HS1 eGFP/mKATE2 fluorescence ratios over 4 hours. Heme synthesis was re-initiated by re-suspending SA-conditioned cells into media lacking SA (black trace). Control cultures that were continuously grown with (green trace) or without (blue trace) SA were used to derive HS1 *R*_max_ and *R*_min_ values, or the fluorescence ratio when the sensor is 10% and 0% bound to heme, respectively. The HS1 fluorescence ratio (*R*) upon the re-initiation of heme synthesis and the *R*_max_ and *R*_min_ values at each time point was entered in to equation 1 (or equation S1), to calculate the percentage of heme bound to HS1, which are presented in **Figures 1c, 1d, 2,** and **3**. All data represent the mean ± SD of triplicate cultures.

**Fig. S3.** A 500 μM dose of succinylacetone (SA) depletes (**a**) total heme and (**b**) heme-loading of the heme sensor, HS1, to values similar to heme deficient *hem1*∆ cells, which lack the 1^st^ enzyme in the heme biosynthetic pathway. All data represent the mean ± SD of triplicate cultures.

**Fig. S4.** Heme sensor expression does not perturb (**a**) the rates of heme synthesis or (**b**) heme trafficking dynamics to the cytosol. (**a**) WT cells expressing *GPD* driven cytosolic (Cyto, black), mitochondrial (Mito, blue), or nuclear (Nuc, green) HS1, or empty vector (EV, grey) were heme depleted with 500 μM succinylacetone (SA) for 15 hours and then the cells were resuspended in fresh SC-LEU media lacking SA, where the re-synthesis of heme was monitored by harvesting 2 x 10^8^ cells every hour and analyzed for heme content as decribed in the **Method** **Details**. (**b**) The heme trafficking dynamics assay was conducted on cells expressing *ADH* (black), *TEF* (green) and *GPD* (blue) driven cytosolic HS1. Despite the ~10-fold increase in HS1 expression between *ADH* and *GPD* promoters, as measured by mKATE2 fluorescence in the right panel, cytosolic heme trafficking rates are virtually identical. All data represent the mean ± SD of triplicate cultures.

**Fig. S5.** Heme binding kinetics of cytosolic, nuclear, and mitochondrial-targeted HS1. 2 x 10^8^ exponential-phase WT cells expressing *GPD* driven cytosolic (Cyto, black), mitochondrial (Mito, blue), or nuclear (Nuc, green) HS1 were lysed in 500 μL of PBS buffer containing 10 mM ascorbate and 0.1% Triton X-100. 100 uL of the cell lysate was analyzed by fluorescence (ex. 488 nm, em. 510 nm; ex. 588 nm, em. 620 nm) over the indicated time period, and at time 0, an automatic dispenser pipetted 5 μL of a 1 mM hemin chloride stock solution in DMSO, giving a final concentration of 50 μM heme in the cell lysate. All data represent the mean ± SD of triplicate cultures.

**Fig. S6.** The effects of *mgm1*∆ and *dnm1*∆ on steady-state HS1 heme loading and heme synthesis. (**a**) Cytosolic (Cyto), nuclear (Nuc), or mitochondrial (Mito) HS1 expressed in WT (black, gray), *dnm1*∆ (dark and light green), and *mgm1*∆ (dark and light blue) cells were cultured for 16 hours in SCE-LEU media with (+SA, dark colors) or without (-SA, light colors) 500 μM succinylacetone (SA). Following growth and washing cells with ultrapure water, HS1 sensor fluorescence was measured in a 100 μL suspension of 5 OD’s/mL in PBS. (**b and c**) The rates of heme synthesis were measured in (**b**) *mgm1*∆ and (**c**) *dnm1*∆ cells by first heme depleting cells with 500 μM SA for 15 hours and then re-initiating heme synthesis by re-suspending cells in media lacking SA. 2 x 10^8^ cells were harvested every hour and analyzed for heme content as described in the **Method** **Details**. All data represent the mean ± SD of triplicate cultures.

**Fig. S7.** Validation of the mitochondrial network morphology defects in yeast fission and fusion mutants. (**a**) Sampling and classification of mitochondrial network morphologies observed using Mitotracker staining of cells. (**b**) Histograms of mitochondrial network morphology in the fission and fusion mutants used throughout this study. Mutants defective in mitochondrial fission exhibit an elongated mitochondrial network. Mutants defective in mitochondrial fusion exhibit a punctate mitochondrial network. WT and *mgm1*∆ *dnm1*∆ cells tend to have a more equal distribution of elongated and punctate mitochondrial networks. The histograms were generated by analyzing ~50 cells per mutant. (**c**) Representative images of the mitochondrial network in the fission and fusion mutants utilized in this study.
