## Supplementary figures and images for "Mitochondrial-nuclear heme trafficking is regulated by GTPases that control mitochondrial dynamics"

### Supplemental Figure 1

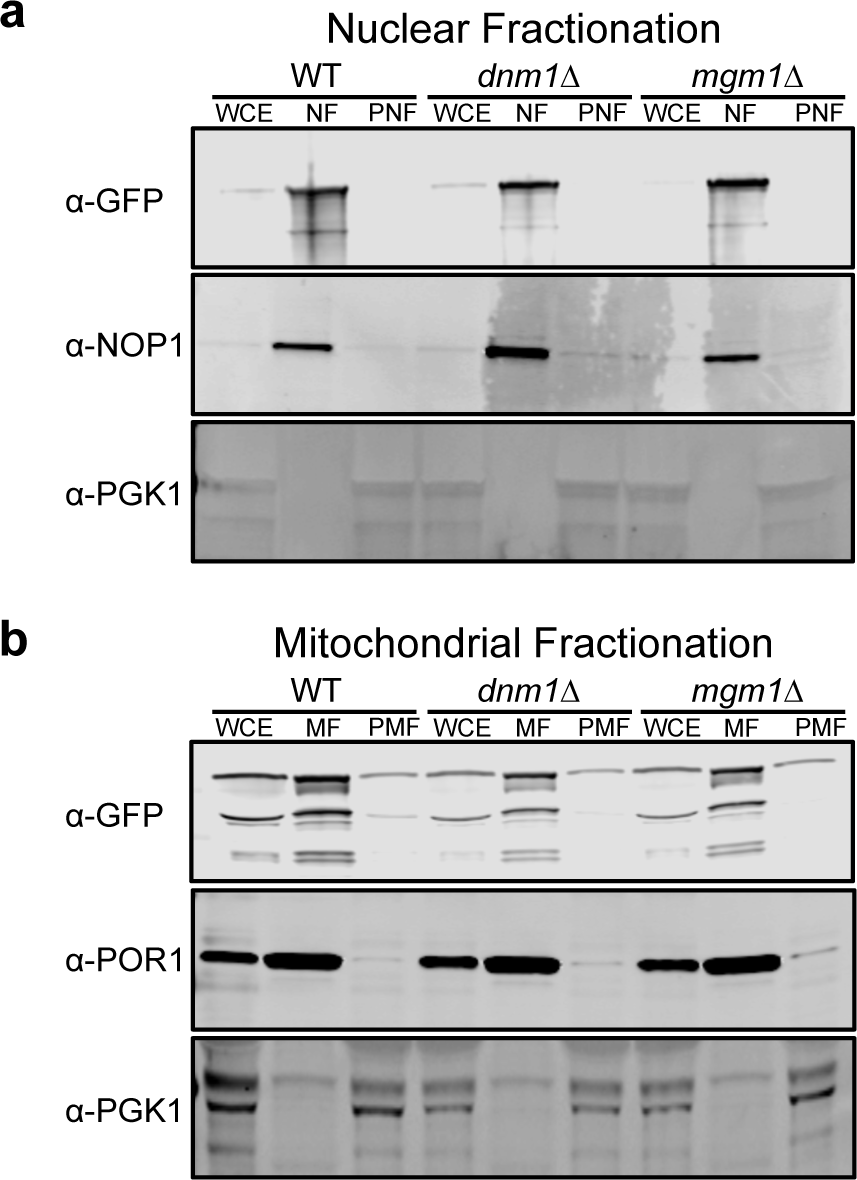

### Supplemental Figure 2

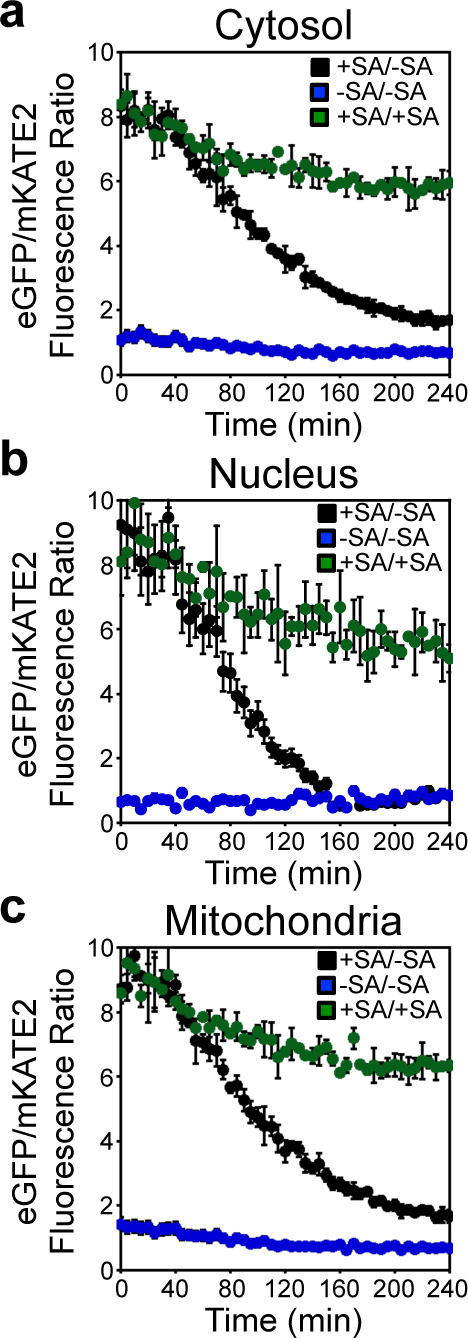

### Supplemental Figure 3

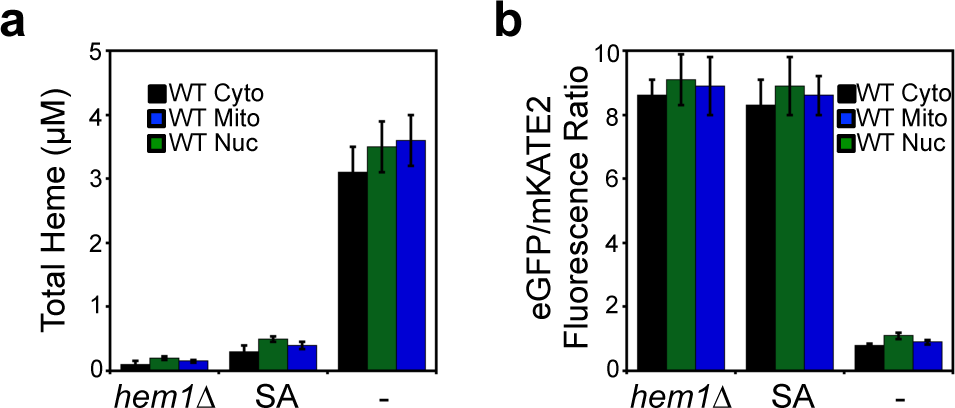

### Supplemental Figure 4

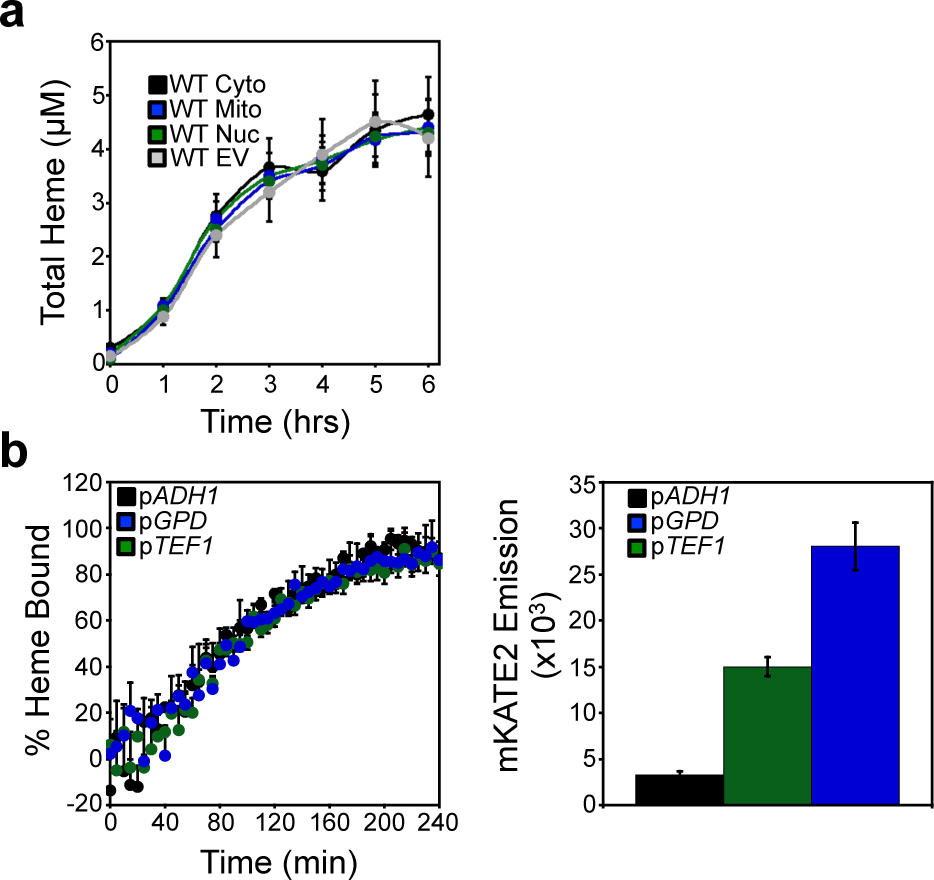

### Supplemental Figure 5

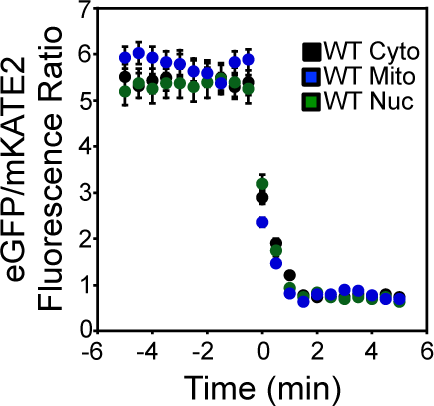

### Supplemental Figure 6

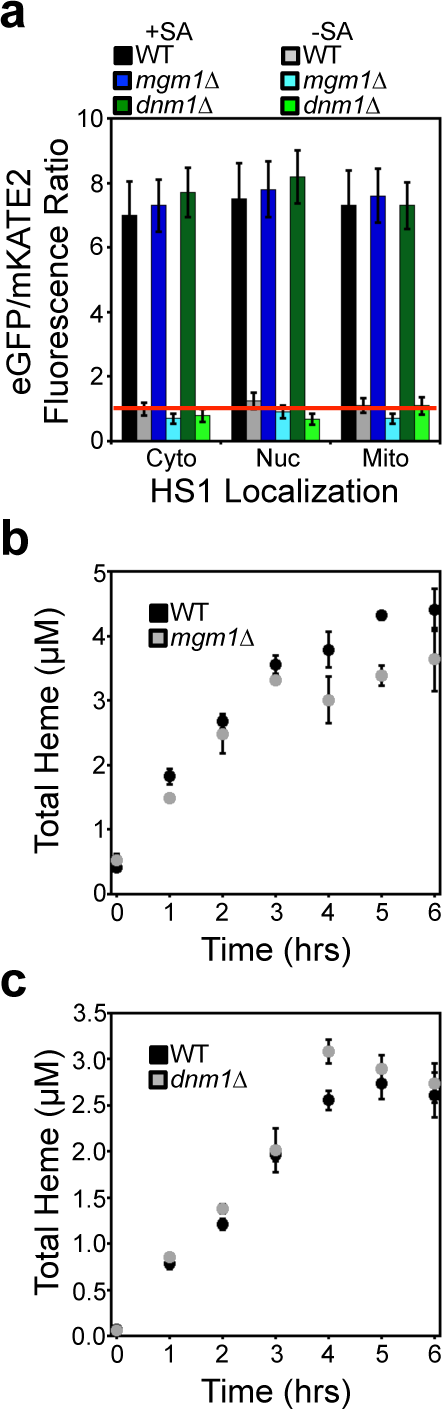

### Supplemental Figure 7

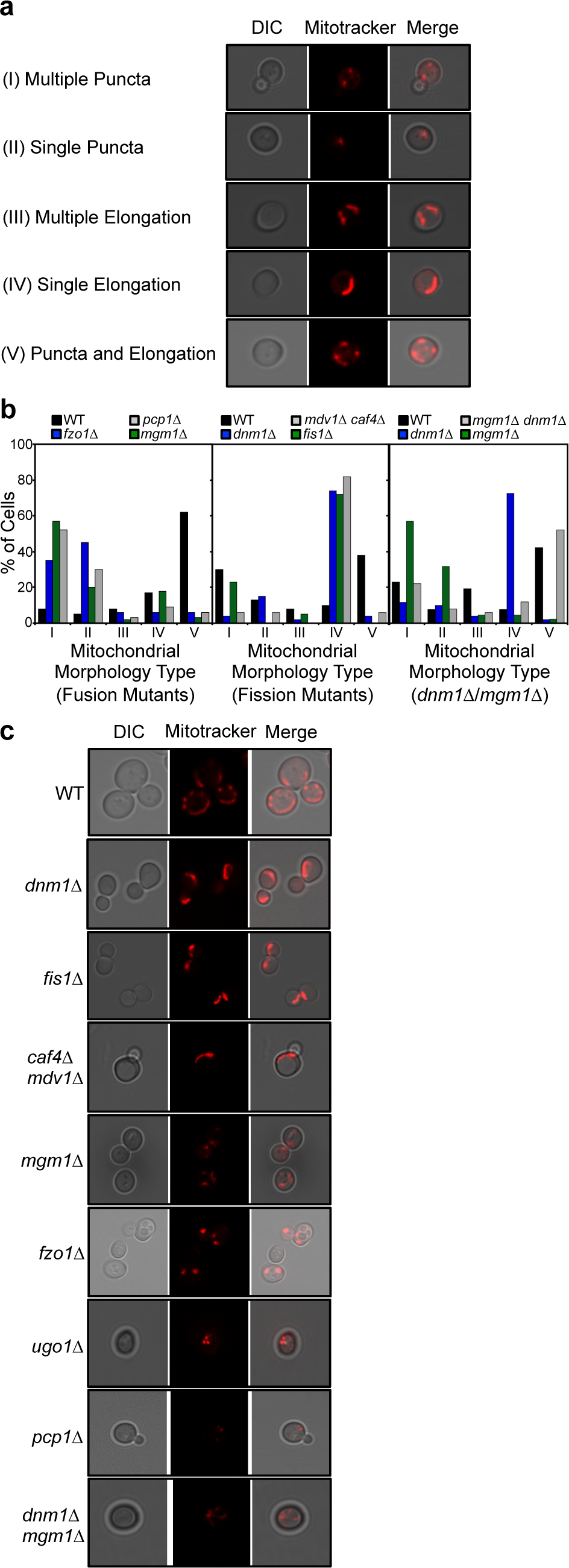
